# basicsynbio and the BASIC SEVA collection: Software and vectors for an established DNA assembly method

**DOI:** 10.1101/2022.01.06.446575

**Authors:** Matthew C. Haines, Benedict Carling, James Marshall, Vasily A. Shenshin, Geoff S. Baldwin, Marko Storch, Paul Freemont

## Abstract

Standardized DNA assembly methods utilizing modular components provide a powerful framework to explore design spaces and iterate through Design-Build-Test-Learn cycles. Biopart Assembly Standard for Idempotent Cloning (BASIC) DNA assembly uses modular parts and linkers, is highly accurate, easy to automate, free for academic and commercial use, while enabling simple hierarchical assemblies through an idempotent format. These features facilitate various applications including pathway engineering, ribosome binding site tuning, fusion protein engineering and multiplexed gRNA expression. In this work we present basicsynbio, an open-source software encompassing a Web App (https://basicsynbio.web.app/) and Python Package (https://github.com/LondonBiofoundry/basicsynbio). With basicsynbio, users can access commonly used BASIC parts and linkers while robustly designing new parts and assemblies with exception handling for common design errors. Users can export sequence data and create build instructions for manual or acoustic liquid-handling platforms. The generation of build instructions relies on the BasicBuild Open Standard which is easily parsed for bespoke workflows and is serializable in Java Script Object Notation for transfer and storage. We demonstrate basicsynbio by assembling a collection of 30 vectors using various sequences including modules from the Standard European Vector Architecture (SEVA). The BASIC SEVA collection is compatible with BASIC and Golden Gate using BsaI. It encompasses vectors containing six antibiotic resistance markers and five origins of replication from different compatibility groups, including a temperature-sensitive variant. To make the collection accessible we deposited it on Addgene under an OpenMTA agreement. Furthermore, vector sequences are accessible from within the basicsynbio application programming interface along with other collections of parts and linkers, providing an ideal environment to design assemblies for bioengineering applications using BASIC.

## Introduction

DNA assembly is an essential tool in Synthetic Biology and Life Sciences, required for building genetic designs and iterating through the Design-Build-Test-Learn cycle^1,2^. A large repertoire of DNA assembly methods are available to researchers and the choice of a suitable method will depend on factors such as the freedom to include forbidden restriction sites, the availability of part libraries or a need for high accuracy^2,3^. Standardized and modular DNA assembly methods are ideal for high-throughput and hierarchical assemblies enabling the costeffective generation of large numbers of constructs with high accuracy while encouraging the reuse of parts across designs^2,4–7^.

BASIC DNA assembly is a standardized DNA assembly method which utilizes modular parts and linkers as functional units^7–10^. The method benefits from several desirable attributes including a single part storage format and assembling up to 14 parts and linkers per round with > 90 % accuracy^7^. Given linkers can encode functional sequences such as ribosome binding sites and fusion protein linkers, diverse constructs are feasible within a single round of assembly. It is also free for academic and commercial use and only requires the absence of one restriction enzyme site (BsaI). It is easy to automate the physical workflow^10^ and conduct hierarchical assemblies since parts are stored in a single format and assemblies are ubiquitously returned with flanking sequences reconstituting this format, enabled by the underlying single-tier, idempotent architecture.

BASIC compares favourably with modular methods based on Golden Gate assembly^5,6^, where multiple restriction enzymes are utilised and assemblies not conforming to standard transcriptional units e.g., operons, are not supported. Notably, BASIC DNA assembly was successfully applied to several areas of Synthetic Biology and Life Sciences research including combinatorial pathway engineering^3,11,12^, synthetic operon^10^ & sRNA circuit design^13^, combinatorial gRNA expression for gene editing^14^, ribosome binding site (RBS) tuning and fusion protein engineering^7^.

In this work, we developed basicsynbio design software, encompassing a Python Package and user-friendly Web App with several aims. Firstly, we make commonly used parts and linkers more accessible *in silico.* Secondly, we prevent assembly and part design mistakes by introducing exception handling. Thirdly, we enable users to export a variety of data types for downstream building, validating, and sharing of assemblies. In particular, we provide users with a data standard describing multiple assemblies which is easily parsed into custom manual and automated workflows. This extends our previous work DNA-BOT^10^, which automated BASIC DNA assembly specifically for the Opentrons platform. We demonstrate basicsynbio by designing and exporting data for a collection of 30 vectors containing several modules from the SEVA database^15,16^. We subsequently build and deposit the collection on Addgene and make the sequences accessible via the basicsynbio API enabling access for BASIC DNA assembly users and the community.

## Materials and methods

### Preparation of BASIC linkers and parts

Apart from BSEVA_L1, all BASIC Linkers were acquired from Biolegio (BBT-18500) and prepped according to the manufacturer’s instructions. Oligos for BSEVA_L1 (Supplementary Table S1) were ordered from Integrated DNA Technologies, Inc. and linker halves prepared as previously described^9^.

Unless specified, all plasmid DNA was prepped using Omega BIO-TEK E.Z.N.A.® Plasmid Mini Kit II according to the manufacturer’s instructions. All plasmid DNA was quantified using Qubit™ dsDNA BR Assay Kit (Thermo Scientific™ Q32850).

Each BASIC SEVA vector is composed of three parts: T0 + marker part, ori + T1 part and mScarlet counter-selection cassette. Initially, each was either chemically synthesised or amplified from SEVA vectors^16^ with primers incorporating *iP* and *iS* sequences upstream and downstream, respectively. The resulting linear sequences were cloned into an appropriate vector, prior to prepping plasmid DNA. Specifically, T0 + marker parts were blunt cloned into pJET1.2 (Thermo Scientific™ K1231) according to the manufacturer’s instructions. ori + T1 and mScarlet counter-selection cassette parts were assembled as described in the supplementary data (oris.gb & addgene_submission.pdf) using BASIC DNA assembly^8^. Constructs were plated on LB-agar (ForMedium) supplemented with 100 μg/mL carbenicillin and incubated at 30 or 37 °C prior to picking colonies and prepping plasmid DNA. All parts were sequence verified via Sanger Sequencing, prior to assembly.

### Assembly and validation of the BASIC SEVA collection

All assemblies were designed *in silico* using basicsynbio (Supplementary Data: addgene_submission.pdf). Echo instructions for the “Assembly reaction” step of the workflow and manual instructions for the entire workflow were exported (see Supplementary Data).

Clip Reaction and Magbead purification steps were implemented as described in the manual instructions (Supplementary Data: BASIC_SEVA_collection_v10_manual.pdf), transferring purified clip reactions to an Echo® Qualified 384-Well Polypropylene Microplate (Beckman 001-14555). Purified clip reactions were mixed by executing the echo_clips_1.csv script (Supplementary Data) on a Beckman Echo 525 Acoustic Liquid Handler, using a 96-well destination plate (Azenta Life Sciences 4ti-0960). ddH2O and 10x assembly buffer solutions were transferred to the same destination plate by executing the echo_water_buffer_1.csv script with both solutions transferred from an Echo® Qualified Reservoir, 2×3 Well, Polypropylene Microplate (Beckman 001-11101). The destination plate containing assemblies was sealed with a PCR foil seal (Azenta Life Sciences 4ti-0550), vortexed and centrifuged prior to incubating at 50 °C for 45 min. 25 μL NEB® 5-alpha Competent *E. coli* cells (C2987) were added to each assembly reaction on ice. Transformation reactions were incubated for 20 min on ice, followed by heat shock at 42 °C for 15 sec, recovery on ice for 2 min, the addition of 150 μL SOC media (ForMedium) and outgrowth at 30 °C for 2 hr. Cells were plated on LB-agar containing antibiotics concentrations illustrated in Figure 4b. Plasmid DNA from pink colonies was prepped as described above for corresponding ori + T1 parts.

Prior to Addgene submission we verified the presence of the correct ori + T1 part, sequencing each vector using the BSEVA_L1_overhang sequencing primer (Supplementary Table S1). The resulting data was analysed using cMatch^17^ to verify homology.

### Availability of sequences

All BASIC SEVA plasmids were deposited on Addgene (Deposit 80391) and made available under Addgene’s OpenMTA agreement.

## Results and Discussion

### basicsynbio workflow

We conceived a typical workflow for users implementing basicsynbio (Figure 1a). Initially users would access collections of parts and linkers available from the basicsynbio API, in addition to importing their own. These BasicPart and BasicLinker objects are combined initiating BasicAssembly objects representing assembled constructs. A key advantage of BASIC DNA assembly is its idempotency, meaning assemblies containing LMP & LMS linkers are themselves BasicParts and can function in subsequently larger constructs. basicsynbio facilitates this, enabling users to convert BasicAssembly objects into BasicParts, ready to initiate next-tier, larger BasicAssembly objects. Once the user has specified all the desired BasicAssembly objects, various data types are available to export (Figure 1a and Supplementary Figure S1). Users can export sequence data representing BasicAssembly and BasicPart objects in GenBank via the Web App or in formats supported by Biopython^18^ via the Python Package. Notably, all features are preserved, maintaining annotations in the resulting assemblies. In addition to exporting sequence data, users can export build instructions, for instance instructions for manual or automated workflows e.g., pdf instructions for manual workflows or csv-files to program a Beckman Echo robot.

**Figure 1.**
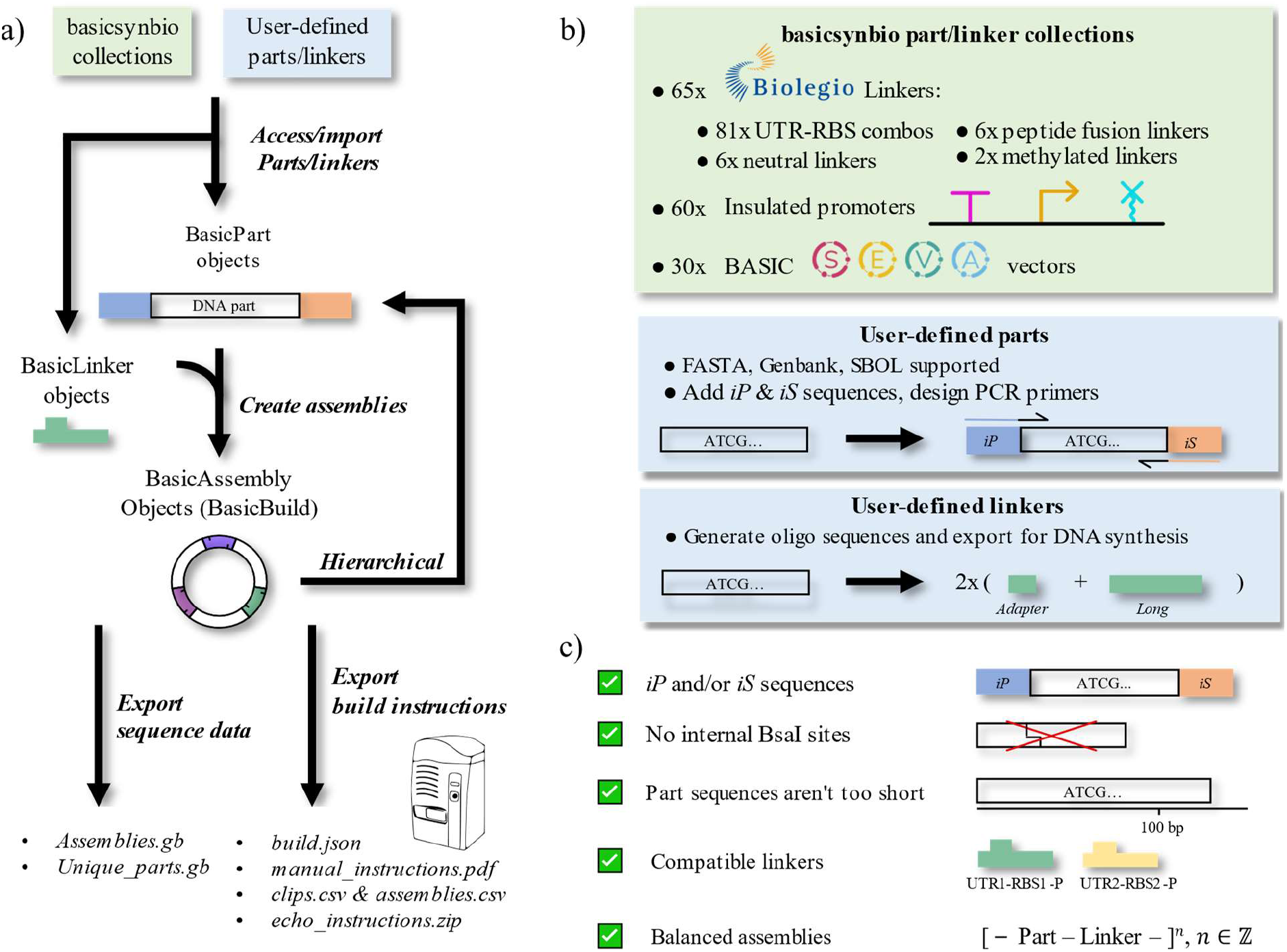
basicsynbio workflow and functionality – a) Typical basicsynbio workflow: BasicPart & BasicLinker objects are imported from internal collections or user-defined sources. Combinations of BasicParts and BasicLinkers initiate BasicAssembly objects, describing assemblies. Sequence data and build instructions are exported for reference and to aid downstream workflows, respectively. Multiple data types are exportable, including those for DNA assembly using an acoustic liquid handler. b) Options for importing parts and linkers. basicsynbio part and linker collections contain > 150 sequences compatible with BASIC DNA assembly, accessible from within the Python Package and Web App. Numerous biological sequence formats are supported for user-defined parts. When creating parts, users can add *iP* and iS sequences and/or design necessary PCR primers. Users can define linkers using the API and export required adapter & long oligonucleotide sequences for prefix and suffix linker halves. c) Error handling in basicsynbio. Objects are checked for common errors that could lead to failure during assembly. For instance, using linker halves incompatible with each other in the same assembly raises an exception, as do unbalanced assemblies.

To aid accessibility of existing core BASIC DNA assembly part and linker sequences, we include PartLinkerCollection objects, accessible from the API and containing instances of commonly used BasicParts and BasicLinkers. Notable collections are illustrated in Figure 1b, including BASIC_BIOLEGIO_LINKERS, BASIC_PROMOTER_PARTS and BASIC_SEVA_PARTS which contain all 65 commercially available Biolegio linkers, including linkers for 81 different untranslated region (UTR)/ribosome binding site (RBS) combinations, a collection of 60 inducible and constitutive promoters, insulated by different combinations of upstream terminators and downstream RiboJ sequences^19^ and a collection of 30 vectors containing several SEVA modules^16^, respectively. To aid the exploration of PartLinkerCollections, users can visualize individual parts and linkers via the Web App using SeqViz or DNAFeaturesViewer^20^ (Supplementary Figure S2 & S3). Different versions of a given PartLinkerCollection are supported enabling future updates where required. Furthermore, users can contribute new PartLinkerCollections as described in the online documentation (https://londonbiofoundry.github.io/basicsynbio/). We hope this will encourage the BASIC DNA assembly user community to share collections of new part and linker sequences for different applications between labs and institutions.

In addition to the above PartLinkerCollections, users can import parts from local files or external sources and/or create new linkers using the API, greatly expanding the number of possible assemblies. Users may import parts specified in commonly used file formats such as FASTA, GenBank and SBOL (Figure 1b & Figure 2). Furthermore, to aid the generation of new parts, users can automatically add required *iP* and *iS* sequences^7^ to the 5’ and 3’ ends of input DNA sequences, respectively. It is also possible to design primers to add iP and iS sequences to parts via overhang PCR with the aid of Primer3^21^. This enables cost-effective conversion of existing DNA sequences into a physical part, avoiding the need for *de novo* DNA synthesis. For a given linker, users can calculate the four oligonucleotide sequences required to generate linker halves, an adapter and long oligonucleotide for each linker half^7,9^ (Figure 1b). This feature aids the generation of custom linkers required for specific applications or organisms.

**Figure 2.**
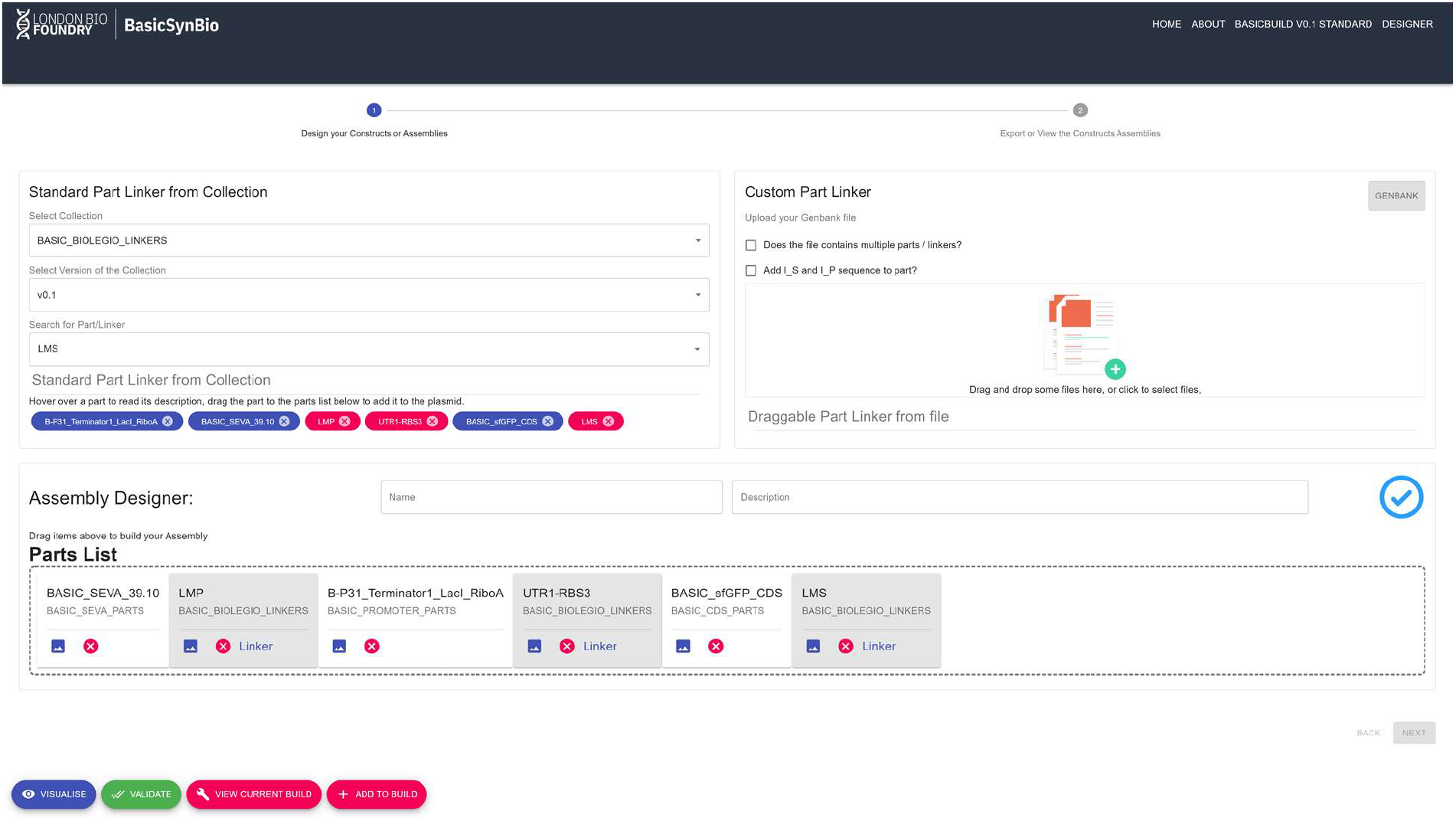
Screenshot of basicsynbio Web App designer. In this example, a construct expressing sfGFP under an IPTG-inducible, insulated promoter is created in the Assembly Designer. BASIC_SEVA_39.10, BP31_Terminator_LacI_RiboA and BASIC_sfGFP_CDS BasicPart objects were selected from BASIC_SEVA_PARTS, BASIC_PROMOTER_PARTS and BASIC_CDS_PARTS collections, respectively. These are combined with LMP, UTR1-RBS3 and LMS linkers from the BASIC_BIOLEGIO_LINKERS collection. The assembly is checked against errors (blue tick) and can be visualized prior to adding to a BasicBuild (bottom-left).

For the successful implementation of BASIC DNA assembly, imported parts and designed assemblies must satisfy several conditions (Figure 1c). For instance, if the length of a part is significantly shorter than 100 bp, the linker-ligated part would be lost during the purification step of assembly. Additionally, internal BsaI sites are not allowed in BASIC parts and specific linkers can only be used once per assembly round, while BasicPart and BasicLinker objects must alternate, with equal numbers of each. Where a user designs an assembly that doesn’t satisfy the above conditions, basicsynbio raises exceptions preventing subsequent experimental failure, increasing robustness.

To implement the basicsynbio workflow illustrated in Figure 1a, users can utilize the open-source Python Package or Web App. Python iterator patterns^22^ combined with the basicsynbio package allow users to initiate large numbers of BasicAssembly objects programmatically, facilitating the exploration of large design spaces with 100s of constructs feasible with BASIC DNA assembly^10^. Meanwhile, the designer interface of the Web App (Figure 2) offers users an intuitive way to create BasicAssembly objects by dragging and dropping selected BasicPart and BasicLinker objects. In addition to visualizing parts using the Web App (Supplementary Figure S2 & S3), users can dynamically visualise assemblies to ensure they contain the desired sequence prior to implementing the checks illustrated in Figure 1c.

### BasicBuild Open Standard

Following design, a user builds their collection of assemblies. To determine build instructions, the user makes multiple calculations^10^. Firstly, the user calculates the unique set of clip reactions required by all assemblies. Each clip reaction is defined by a BasicPart in combination with BasicLinker prefix and suffix halves. Secondly, the user needs to associate each unique clip reaction with the assemblies requiring it. From this, the user calculates the absolute number of each clip given each can support 15-30 assemblies, depending on the workflow. Thirdly, the user makes calculations ensuring a final part concentration of 2.5 nM following clip reaction setup, maximising efficiency. These three parameters guide liquid-handling operations during clip reaction setup and assembly stages of the BASIC workflow. Previously, we implemented this for a specific liquid-handling platform^10^ and in this work we describe a standard enabling bespoke manual and automated workflows.

The BasicBuild Open Standard is a data structure given in JavaScript Object Notation (JSON) which contains the information outlined above. Figure 3 illustrates the four nested objects from an example. The clips_data object contains data on each clip reaction required for the build. Further information on each component is available within unique_parts and unique_linkers objects, where the corresponding key can be used to access this information e.g., “UP0” to access the first part within unique_parts. A link between each clip reaction and the assemblies using it is established by both the “assembly keys” attribute in the clips_data object and the “clips reactions” attribute of the assembly data object. The absolute number of each clip reaction can be calculated using the clips_data “total assemblies” attribute, considering the number of assemblies supplied by each purified clip reaction (15 – 30 depending on the method). To aid the addition of parts to a final concentration of 2.5 nM in clip reactions, a “Part mass for 30 μL clip reaction (ng)” attribute is provided, where the addition of the associated mass to a 30 μL clip reaction results in the desired final concentration. We implement the BasicBuild Open Standard within the basicsynbio API as a python class. To demonstrate the standard’s flexibility, we have written functions that parse BasicBuild class objects into manual instructions in pdf format and instructions for a Beckman Echo liquid handling platform.

**Figure 3.**
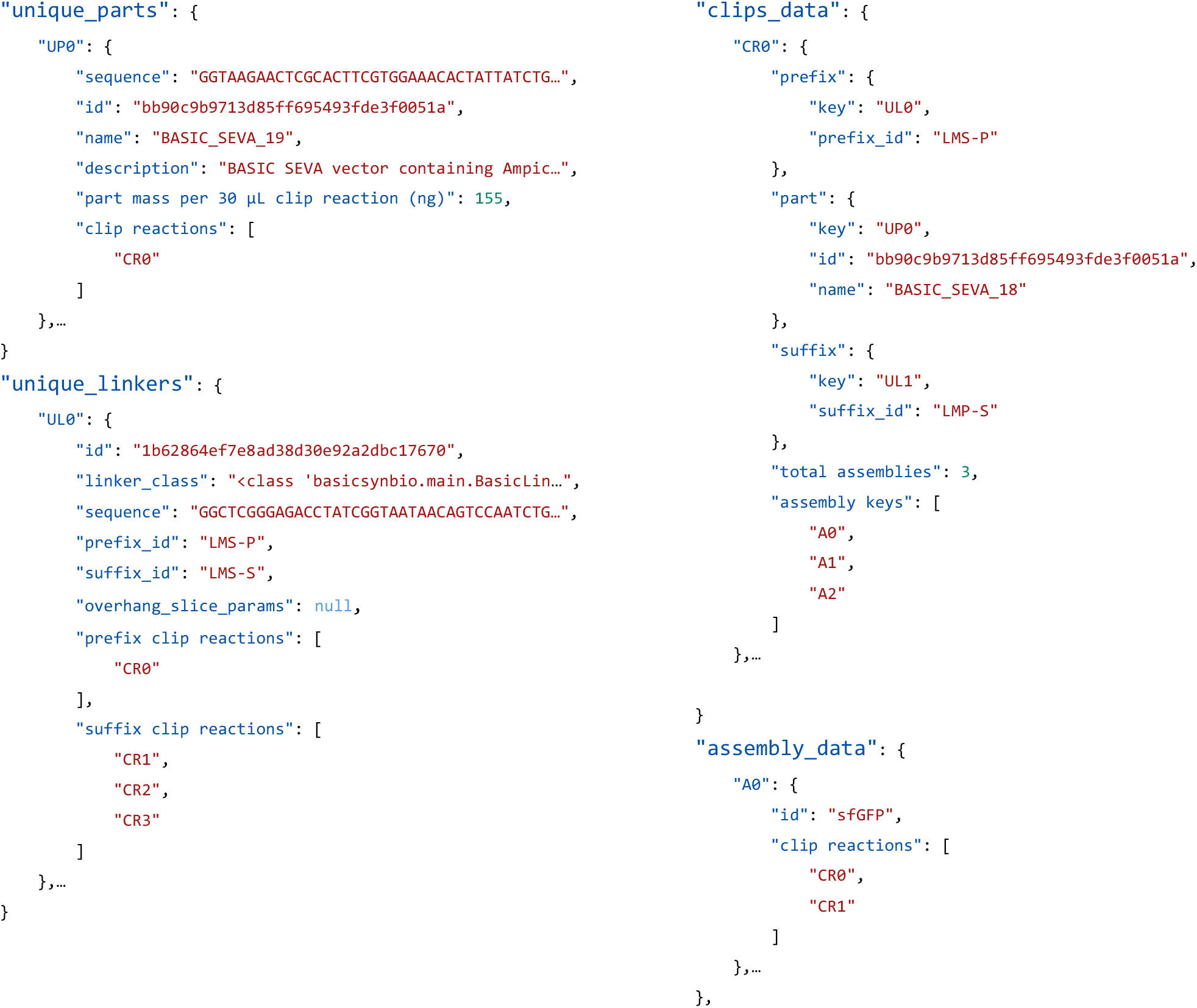
BasicBuild Open Standard – Key objects from a BasicBuild example, serialized in JSON: “unique_parts”, “unique_linkers”, “clips_data” and “assembly_data”. For clarity values have been shortened and similar objects replaced with “…”. Using the BasicBuild Open Standard, the generation of functions for custom workflows is simplified as demonstrated in this work for manual and acoustic liquid-handler instructions. A specification of the BasicBuild Open Standard v0.1 is available online (https://basicsynbio.web.app/basicbuild-standard).

It is also worth noting that the basicsynbio API can decode BasicBuilds serialized in JSON creating a class object. As such, designs serialized in one location can be transferred securely to the location of manufacturing, decoded, and processed into build instructions specific to the facility. We envision this will allow designers to work agnostically of the protocol or facility used for building, freeing them to focus on other steps of the Design-Build-Test-Learn cycle.

### BASIC SEVA collection

To demonstrate basicsynbio, we designed a collection of vectors illustrated in Figure 4a. Each contains a specific combination of antibiotic resistance marker and origin of replication (ori) flanked by SEVA T0 and T1 terminators. The terminators prevent transcriptional readthrough, maintaining plasmid stability in an equivalent manner to vectors from the SEVA database^15^. To enable these vectors to function in BASIC DNA assembly and Golden Gate workflows using BsaI, we flank the terminator, resistance marker and ori components with LMP and LMS linkers. In contrast to vectors from the SEVA database we omit the origin of transfer (OriT) sequence from our collection. This reduces the risk of unintended transfer to natural microorganisms and if desired, users can incorporate this sequence when generating constructs for specific applications.

**Figure 4.**
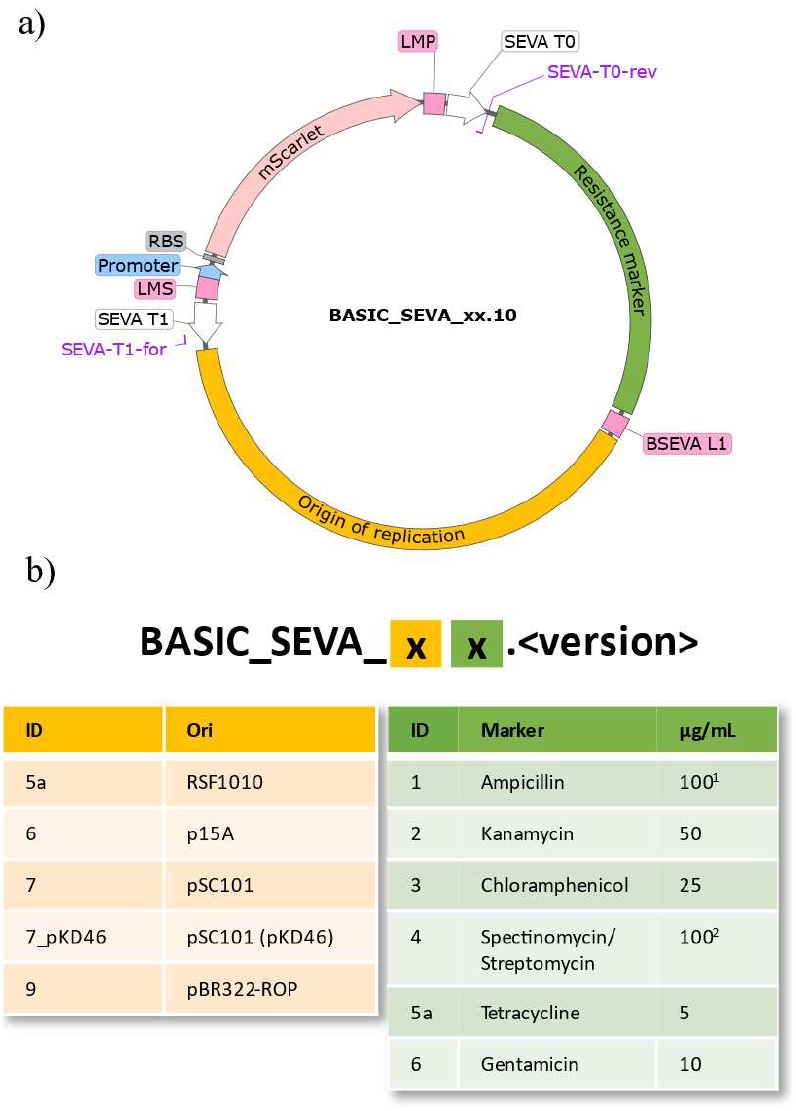
BASIC SEVA vector collection – a) Plasmid map of the collection. BASIC linkers, and SEVA terminators are indicated by magenta and white features, respectively. Origins of replication (yellow) and antibiotic resistance markers (green) are joined via a neutral “BSEVA L1” linker. An mScarlet counter-selection cassette, compromising promoter (blue), RBS (grey) and mScarlet CDS is flanked by methylated LMP and LMS linkers, resulting in drop-out during assembly. SEVA-T0-rev & SEVA-T1-for sequencing primer binding sites are indicated. b) Nomenclature of the BASIC SEVA collection. Vectors are named through the combination of two strings that reflect the origin of replication and resistance marker identities. Where an integer value is given the associated module is identical to that provided by SEVA^16^. The remaining modules are discussed in the text. The nomenclature also provides a versioning system, equivalent to that used by GenBank (https://www.ncbi.nlm.nih.gov/genbank/release/). Antibiotic concentrations used to isolate plasmids in this study are given (^1^carbenicillin was used instead of ampicillin; ^2^streptomycin was not tested in this study but has previously been demonstrated^32^).

The marker and ori are joined with a neutral BASIC linker sequence (BSEVA L1), enabling combinatorial assembly of the collection using BASIC. This linker sequence was designed using DNA Chisel^23^ to retain maximum plasmid stability. Specifically, sequences that could lead to expression such as promoters and RBSs were avoided and complementarity to the *E. coli* MG1655 genome or existing BASIC parts & linkers was minimized, reducing the propensity for recombination in strains with active homologous recombination machinery^24^.

Since the sequence outside the LMP/S flanked region is lost during the assembly of downstream constructs using these vectors, we incorporated an mScarlet counter-selection cassette within this region. Thus, colonies containing these vectors display a pink phenotype allowing for visual counterselection against assembly background from undigested, original vector.

The BASIC SEVA collection generated in this work encompasses 30 vectors, every combination of six markers and five oris described in Figure 4b. The six markers correspond with the first six markers used by the SEVA collection. Apart from the tetracycline module, all marker modules are identical in sequence to SEVA modules. For reasons outlined in the Supplementary Information (BASIC SEVA modules) we use an alternative tetracycline sequence but note its difference to that used by SEVA by assigning a “5a” ID as opposed to “5”. We selected four oris from the SEVA database to include in our collection with three identical in sequence (6, 7 & 9). The fourth (ori 5a) is homologous to the SEVA RSF1010 module with the sequence previously reported in the literature^25^.

To assemble the collection, we split the design in Figure 4a into three BASIC parts: T0 + marker part, ori + T1 part and mScarlet counterselection cassette. Prior to assembling the collection, we cloned, prepped and sequence verified each part (Materials and methods). We subsequently assembled each vector of the collection *in silico* using basicsynbio (Supplementary Data: addgene_submission.pdf), naming each vector according to the BASIC SEVA nomenclature (Figure 4b). Using the resulting BasicBuild we exported manual instructions for the entire assembly and Echo liquid-handling instructions for the “Assembly reaction” step of the workflow, aiding building (Materials and methods). Following assembly and transformation, we selected colonies containing the antibiotic and counter-selection cassette using relevant antibiotics (Figure 4b) and by picking pink colonies, respectively.

Sequencing the collection, we identified deletion or insertion mutations at the border of the gentamicin CDS N-terminus and 5 ‘-UTR for plasmids containing p15A and pBR322 oris (BASIC_SEVA_66.10 & 69.10). We isolated plasmid DNA from a further three colonies for each design and observed similar mutations (Supplementary Figure S6). In contrast to pSC101 and RSF101, both p15A and pBR322 belong to the ColE1 class of plasmids^26^ where copy number is regulated by RNA transcripts^27^. It is conceivable that the SEVA gentamicin cassette sequence interferes in this process though this remains to be determined.

To provide users with plasmids conferring gentamicin resistance and containing p15A & pBR322 oris, we selected constructs from those sequenced having identical mutations and designate them BASIC_SEVA_66.11 & 69.11 (Supplementary Figure S6). A search revealed this mutation in the gentamicin cassette has previously been reported (AJ247370.1). Sequence data describing the entire collection was exported with the assistance of basicsynbio and used to generate a basicsynbio PartLinkerCollection making the BASIC SEVA collection accessible to other basicsynbio users via the API.

The 5 ori modules used to build the collection enable a variety of applications. Notably, we include a temperature-sensitive ori (7_pKD46), not present in the SEVA database. This ori is identical in sequence to the ori present in pKD46^28^. Plasmids harbouring this ori are ideal for applications where plasmid curing is a requirement e.g. strain engineering^28–30^. The three SEVA oris used (6, 7 & 9) are from three different incompatibility groups^31^, enabling applications requiring multiple plasmid types in the same host. Furthermore, these three oris provide a range of copy numbers to tune gene expression. As previously discussed^15^, the remaining ori (RSF1010) is compatible with a broad range of hosts enabling applications suited to non-model organisms. While a high copy number ori was not included in the collection (Supplementary information - BASIC SEVA modules), plasmids containing pBR322 can be amplified with the addition of chloramphenicol, increasing plasmid yield^31^. Additionally, we observed a relatively higher yield for vector BASIC_SEVA_39.10 which contains both pBR322 ori and a chloramphenicol marker (data not shown), suggesting this vector is suitable for applications requiring high yields of plasmid DNA.

In conclusion, with the basicsynbio Python Package and Web App users can access commonly used parts and linkers, robustly design new parts, linkers, and assemblies while exporting sequence data and build instructions. We also outline the BasicBuild Open Standard, enabling the facile generation of custom build instructions as demonstrated in this work for manual workflows and workflows using acoustic liquid-handlers. To demonstrate basicsynbio we design and assemble a collection of 30 vectors using modules from the SEVA database. Sequence data for this collection is available for users via the basicsynbio API and plasmids were deposited on Addgene. In combination with other accessible parts and linkers users can easily and robustly design a large repertoire of assemblies enabling applications in Synthetic Biology and the Life Sciences.

## Author Contributions (CRediT author statement)

M.C.H.: Conceptualization, Investigation, Software, Visualization, Writing – Original Draft, Writing – Review & Editing, Project Administration

B.C.: Software, Writing – Review & Editing, Visualization

J.M.: Investigation, Writing – Review & Editing

V.A.S.: Investigation, Writing – Review & Editing

G.S.B.: Conceptualization, Supervision, Writing – Review & Editing

M.S.: Conceptualization, Investigation, Supervision, Project

Administration, Writing – Review & Editing

P.F.: Funding Acquisition, Supervision, Writing - Review & Editing.

## Acknowledgments

P.F. and M.C.H. are supported by UKRI Engineering and Physical Sciences Research Council (EP/T013788/1). We also thank Alexis Casas for discussion on software and the BasicBuild Open Standard.

