## Supplementary_Information for "basicsynbio and the BASIC SEVA collection: Software and vectors for an established DNA assembly method"

#### BASIC SEVA modules

As outlined in the main text, we utilised marker 5a sequence over the SEVA tetracycline module and were unable to generate BASIC SEVA plasmids containing a high copy number origin of replication. The following sections provide an overview of these findings.

##### Marker 5a

After cloning the SEVA tetracycline module into pJET1.2 (Thermo Scientific™ K1231), we identified a 248I>S substitution in the amino acid sequence of the tetracycline efflux transporter, we assigned this variant the identity BS_5axx. We conducted an NCBI BLASTp search against the amino acid sequence but failed to find a variant containing the 248I>S substitution.

We attempted to correct this error via PCR mutagenesis, amplifying the module cloned into pJet1.2 (Thermo Scientific™ K1231) with the BS_5xx_980G>T_F/R primer pair (Table 1) and plating cells transformed with the circularised product on LB-agar supplemented with 100 µg/mL carbenicillin. The equivalent SEVA module sequence (BS_5xx) was confirmed via Sanger Sequencing.

We measured the transformation efficiency (TE) of BS_5xx and compared it to BS_5axx (Figure S4). We found BS_5axx had a larger TE compared to BS_5xx ($P < 0.05$) with mean TEs of $1.8 \times 10^7$ and $1.3 \times 10^5$, respectively. The reason for the difference in TE was not investigated further but could indicate BS_5xx does not efficiently confer tetracycline resistance under the conditions tested, requiring additional mutations. We therefore opted to use BS_5axx when generating the BASIC SEVA collection described in this paper.

##### pUC origin of replication

We attempted to include the SEVA pUC origin of replication module (SEVA #8) in the BASIC SEVA collection. In a similar manner to that conducted for other BASIC SEVA oris (Materials and methods), we assembled the SEVA #8 ori + T1 part with a T0 + ampicillin marker part, a construct we refer to as BS_x8x. Following plasmid DNA preparation and sequencing, we identified a 1089T>A point mutation in the resulting BS_x8x1089T>A plasmid. We attempted to correct his error via PCR mutagenesis using the BS_x8x1089A>T_F/R primer pair (Table 1). We transformed *E. coli* DH5α cells with both the circularised and linear PCR mutagenesis products (Figure S5). Transformation of linear PCR products gives an indication of background, a result of BS_x8x1089A>T PCR template not digested by a subsequent DpnI step. We observed many microcolonies following transformation with the circularised PCR product, an indication of a burdensome construct (Figure S5). We also observed some larger colonies, though not many more than achieved via transformation of linear PCR product. We prepped plasmid DNA from two of the larger colonies, however both contained further mutations to the pUC origin of replication (data not shown). We conclude BS_x8x is too burdensome for DH5α. As discussed in the main text, we recommend the use of BASIC_SEVA_39.10 for applications requiring a high yield of plasmid DNA or where another marker is required, we suggest amplifying a plasmid containing a pBR322 ori with chloramphenicol[1].

### Supplementary Figures

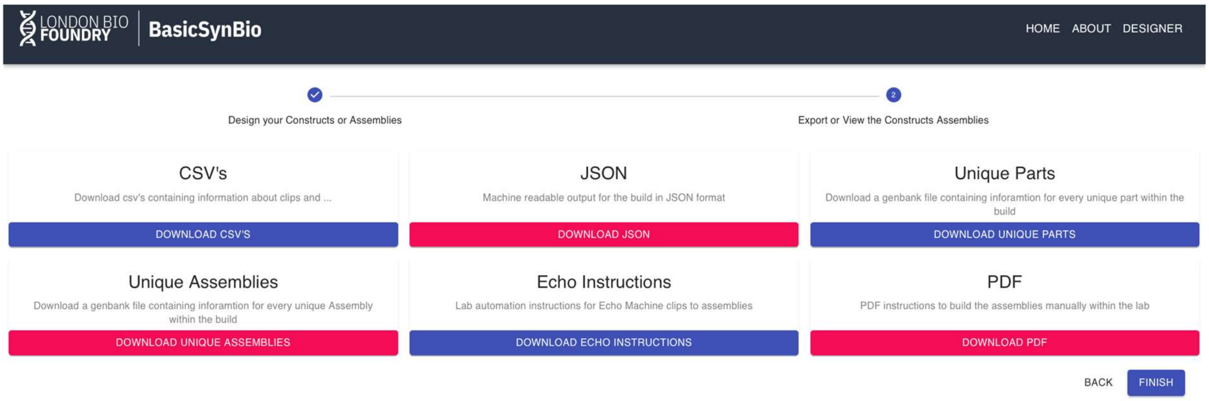

**Figure S1. A screenshot displaying the export options available to basicsynbio Web-app users.**

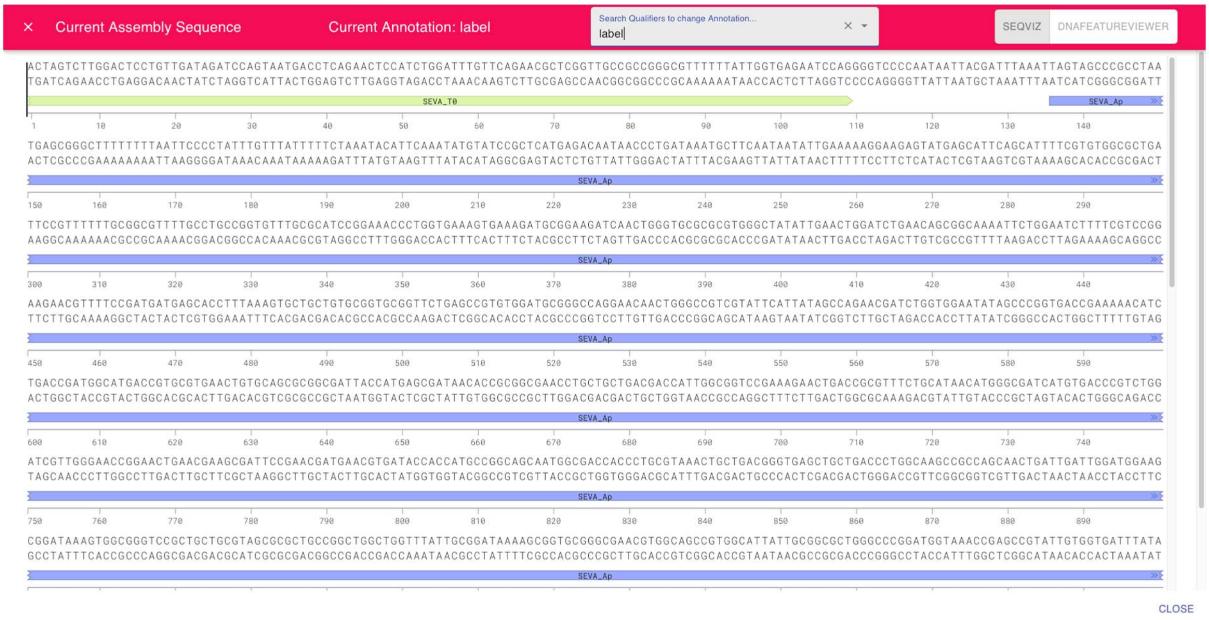

**Figure S2. A screenshot highlighting the integration of Seqviz within the basicsynbio Web-app which can be used to view the sequence of any BasicPart or BasicAssembly.**

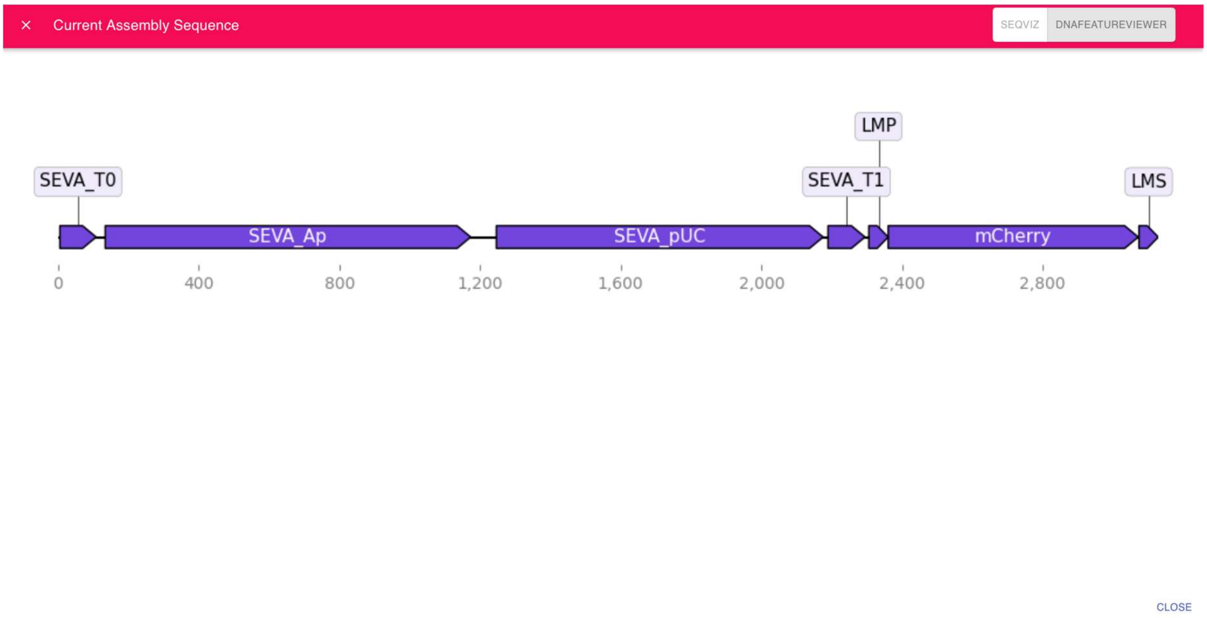

**Figure S3. A screenshot highlighting the integration of DnaFeaturesViewer within the basicsynbio Web-app which can be used to view the sequence of any BasicPart or BasicAssembly.**

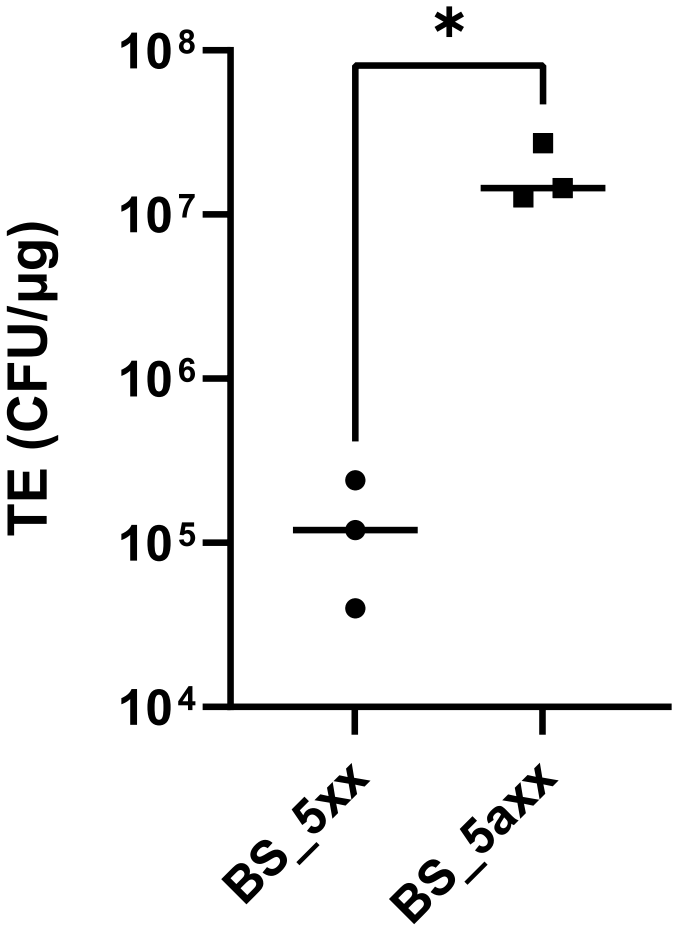

**Figure S4. Scatter plot illustrating transformation efficiency (TE) of considered tetracycline marker modules for the BASIC SEVA collection as outlined in section "Marker 5a". TE was measured by calculating colony forming units (CFU) per µg DNA transformed into chemically competent cells (NEB C2987), as described by the manufacturer, when reactions were plated on LB-agar supplemented with 10 µg/mL tetracycline. Mean TE is illustrated by a horizontal line. Significance of the difference was measured using Welch's t-test (One-tail), where * denotes P<0.05.**

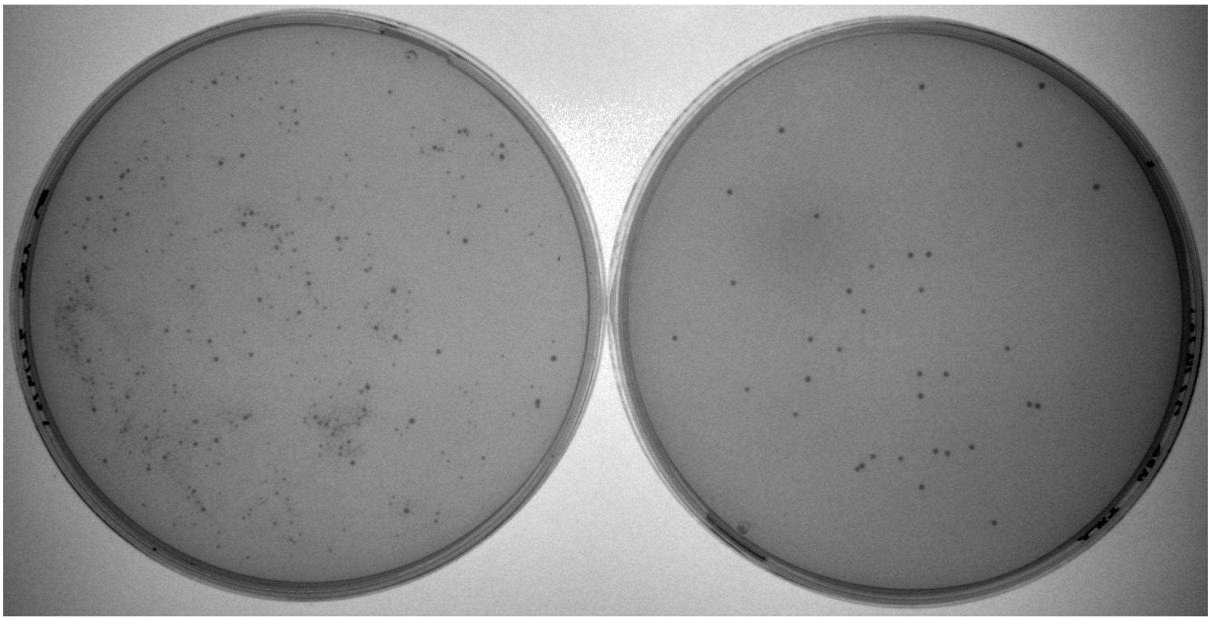

**Figure S5. PCR mutagenesis of BS_x8x1089T>A. Left: *E. coli* DH5α transformed with circularised product of PCR mutagenesis (putative BS_x8x). Right: *E. coli* DH5α transformed with linear product of PCR mutagenesis.**

```
SEVA_Gm                      -------------------AGCAGCAACGATGTTACGCAGCAGCAACGATGTTACGCAGC
BASIC_SEVA_66_clone_E11      --------------------------------------AGCAGCAACGATGTTACGCAGC
BASIC_SEVA_66_clone_F11      --------------------------------------AGCAGCAACGATGTTACGCAGC
BASIC_SEVA_66_clone_G11      AGCAGCAACGATGTTACGCAGCAGCAACGATGTTACGCAGCAGCAACGATGTTACGCAGC
BASIC_SEVA_66_clone_H11      AGCAGCAACGATGTTACGCAGCAGCAACGATGTTACGCAGCAGCAACGATGTTACGCAGC
BASIC_SEVA_69_clone_E12      --------------------------------------AGCAGCAACGATGTTACGCAGC
BASIC_SEVA_69_clone_F12      --------------------------------------AGCAGCAACGATGTTACGCAGC
BASIC_SEVA_69_clone_G12      AGCAGCAACGATGTTACGCAGCAGCAACGATGTTACGCAGCAGCAACGATGTTACGCAGC
BASIC_SEVA_69_clone_H12      AGCAGCAACGATGTTACGCAGCAGCAACGATGTTACGCAGCAGCAACGATGTTACGCAGC
```

**Figure S6. Multiple sequence alignment of the gentamicin resistance cassette region from BASIC_SEVA_66 & 69 clones. The sequence of the original SEVA_Gm part is given. The predicted ATG start codon of the gentamicin resistance CDS is highlighted by a light blue rectangle. Clones H11 & G12 were designated BASIC_SEVA_66.11 & 69.11, respectively.**

#### Supplementary Tables

**Table 1. Oligonucleotide used in this work. "/5Phos/" denotes the presence of a 5′ phosphate group.**

| Primer name | Oligonucleotide sequence | Comment |
|---|---|---|
| **BS_5xx_980G>T_R** | /5Phos/GCTCCAGCGAAAGCGGTC | Mutagenesis primers |
| **BS_5xx_980G>T_F** | /5Phos/GCGACGATGAtCGGCCTGTCG | |
| **BS_x8x1089A>T_R** | /5Phos/GTAGGTATCTCAGTTCGG | |
| **BS_x8x1089A>T_F** | /5Phos/AGCGTGAGCTtTGAGAAAGCG | |
| **SPAP** | CGAGCCGTATTGTGGTGATTTA | Sequencing primers |
| **SP072** | GGACCCCTGGATTCTCACC | |
| **pJET1.2 Forward** | CGACTCACTATAGGGAGAGCGGC | |
| **pJET1.2 Reverse** | AAGAACATCGATTTTCCATGGCAG | |
| **BS_x5x 1** | AAAGGCGCTCGATGCAG | Additional sequencing primers to sequence BS_x5ax ori + T1 part |
| **BS_x5x 2** | ACACGATAGAGCACCCGG | |
| **BS_x5x 3** | AAAGGCTTGTCTTCGCGG | |
| **BS_x5x 4** | ATTCACGCAGCAGCACC | |
| **BSEVA_L1_prefix_long** | /5Phos/GGACACTATCTGTGTGGACCTCAGAATTGTACGAGAC | BSEVA_L1 oligos. All ordered HPLC-purified. |
| **BSEVA_L1_prefix_adapter** | /5Phos/CACACAGATAGT | |
| **BSEVA_L1_suffix_long** | /5Phos/CTCGGCCCACTTGTGTGTCTCGTACAATTCTGAGGTC | |
| **BSEVA_L1_suffix_adapter** | /5Phos/ACACAAGTGGGC | |
| **BSEVA_L1_overhang** | GTCTCGTACAATTCTGAGGTC | |
