## Supplementary_Data for "basicsynbio and the BASIC SEVA collection: Software and vectors for an established DNA assembly method": addgene_submission_notebook.pdf

### basicsynbio Addgene submission collection

This Notebook contains code to aid the generation and submission of BASIC SEVA plasmids to addgene.

#### Notes on preperation

- [x] The initial p15A ori was corrected given it had a "t" insertion, yielding `BS_x6x`.
- [x] The Benchling feature library (BASIC_SEVA_benchling_misc_features_library.csv) was updated commit: `a2fb18c`.

#### Aims and objectives for cell/s below

- [x] Import the following objects:
  - [x] pickled BasicLinker `BSEVA_L1`
  - [x] Selected Oris and AbR markers specified in #130.
  - [x] mScarlet counter selection casettes for associated Oris.
- [x] Assemble BS_x7x_TS gblocks with BS_1xx to place them in storage vectors. Export associated data.
- [x] Generate a dictionary mapping between Oris and counter-selection casettes.
- [x] Make assemblies and export the following data:
  - [x] Genbank file containing all correctly assembled backbones.
  - [x] Manual and automated build instructions.
- [x] Calculate mass of AbR markers to add to clip reaction. *pJet contains multiple BsaI sites and hence cannot be imported*.

#### Notes on implementation of above

- Following commit `4378e7e` x9x constructs were assembled with the 405 counter-selection cassette instead of the 407 cassette. This was in an attempt to lower the burden of these constructs.

In [ ]:
```python
import basicsynbio as bsb
from basicsynbio.cam import seqrecord_hexdigest
from basicsynbio.utils import MARKER_DICT, ORI_DICT
from Bio import SeqIO
import csv
import numpy as np
import pandas as pd
from pathlib import Path
import pickle
from platemap import Plate
import re
from typing import Tuple, Dict
```

In [ ]:
```python
PATH_TO_SEQS = Path.cwd().parents[0] / "sequences"
PATH_TO_INITIAL_MODS = PATH_TO_SEQS / "genbank_files" / "BASIC_SEVA_collection" / "initial_mods"
# Import objects
linkers = bsb.BASIC_BIOLEGIO_LINKERS["v0.1"]
with open(PATH_TO_SEQS / "alternative_formats" / "pickles" / "SEVA-BB1", "rb") as file:
    bb_linker = pickle.load(file)
abr_markers = list(
    bsb.import_parts(PATH_TO_INITIAL_MODS / "AbR_markers.gb",
    "genbank"
))
oris = list(
    bsb.import_parts(PATH_TO_INITIAL_MODS / "oris.gb",
    "genbank"
)) # collection of oris previously assembled with BS_1xx
cs_cassettes = list(
    bsb.import_parts(PATH_TO_INITIAL_MODS / "mScarlet_cs_cassettes.gb",
    "genbank"
))
ts_ori_gblocks = list(bsb.import_parts(PATH_TO_INITIAL_MODS / "TS_ori_gblocks.gb", "genbank"
))
# Check for unwanted Benchling annotations
initial_modules = abr_markers + oris + cs_cassettes
unwanted_annotations = ("Translation", "Primer")
for initial_module in initial_modules:
    if initial_module.id == "BS_5axx":
        continue
    qualifier_values = [value[0] for feature in initial_module.features for value in list(feature.qualifiers.values())]
    for unwanted_annotation in unwanted_annotations:
        re_matches = [re.match(unwanted_annotation, value[:len(unwanted_annotation)]) for value in qualifier_values]
        if re_matches.count(None) != len(re_matches):
            raise ValueError(f"{initial_module.id} contains unwanted annotations")
```

In [ ]:
```python
bs_1xx = [marker for marker in abr_markers if marker.id == "BS_1xx"][0]
# Assemble BS_x7x_TS
ts_ori_assemblies = [bsb.BasicAssembly(
    ts_ori.id,
    linkers["LMP"],
    ts_ori,
    linkers["LMS"],
    bs_1xx
) for ts_ori in ts_ori_gblocks]
bsb.export_sequences_to_file(
    ts_ori_assemblies,
    PATH_TO_INITIAL_MODS / "TS_ori_modules.gb"
)
bsb.pdf_instructions(
    bsb.BasicBuild(*ts_ori_assemblies),
    "/home/hainesm6/github_repos/LondonBiofoundry/basicsynbio/pdfs/bs_x7x_ts_oris_instructions.pdf",
)
# Assemble counter-selection cassettes
cs_cassette_ori_mapping = {
    "B405_J23106-RBS34-mScarl": "BS_x9x",
    "B407_J23119-RBS34-mScarl": "BS_x6x",
    "B408_J23119-RBS-A12-mSc": "BS_x7x",
}
cs_cassettes = [
    bsb.BasicAssembly(
        cs_cassette.id,
        linkers["LMP"],
        cs_cassette,
        linkers["LMS"],
        bs_1xx,
        bsb.BASIC_BIOLEGIO_LINKERS["v0.1"]["L1"],
        [ori for ori in oris if ori.id == cs_cassette_ori_mapping[cs_cassette.id]][0]
    ).return_part(
        id=cs_cassette.id,
        name=cs_cassette.id,
    ) for cs_cassette in cs_cassettes
]
```

```
/home/hainesm6/.pyenv/versions/basicsynbio/lib/python3.8/site-packages/Bio/SeqIO/InsdcIO.py:726: BiopythonWarning: Increasing length of locus line to allow long name. This will result in fields that are not in usual positions.
  warnings.warn(
```

In [ ]:
```python
def bseva_assembly(
    marker: bsb.BasicPart,
    ori: bsb.BasicPart,
    cs_cassette: bsb.BasicPart,
    id: str ="foobar",
) -> bsb.BasicAssembly:
    """Return a BASIC SEVA backbone as an assembly object.

    Args:
        marker: Antibiotic resistance marker.
        ori: Origin of replication.
        cs_cassette: Counter-selection containing mScarlet CDS.
        id: ID for assembly
    """
    return bsb.BasicAssembly(
        id,
        linkers["LMP"],
        marker,
        bb_linker,
        ori,
        linkers["LMS"],
        cs_cassette
    )


def marker_abbreviation(marker_id: str) -> str:
    """Return abbreviation for resistance marker."""
    if re.match("BS_\daxx", marker_id) != None:
        return abr_marker.id[-4:-2]
    return abr_marker.id[-3]


# Map oris with counter selection cassettes
ori_cs_cassette_mapping = {
    "BS_x5ax": "B407_J23119-RBS34-mScarl",
    "BS_x6x": "B407_J23119-RBS34-mScarl",
    "BS_x7x": "B408_J23119-RBS-A12-mSc",
    "BS_x7x_pKD46": "B408_J23119-RBS-A12-mSc",
    "BS_x9x": "B405_J23106-RBS34-mScarl",
}
ori_cs_cassette_mapping = {
    key: cs_cassettes[[cs_cassette.id for cs_cassette in cs_cassettes].index(value)] for key, value in ori_cs_cassette_mapping.items()
}
# Make assemblies and export data
MARKER_DICT = MARKER_DICT.copy()
MARKER_DICT["5a"] = "Tetracycline"
assemblies = []
for abr_marker in abr_markers:
    abr_abbreviation = marker_abbreviation(abr_marker.id)
    for ori in oris:
        assembly = bseva_assembly(
            abr_marker,
            ori,
            ori_cs_cassette_mapping[ori.id]
        )
        assembly.marker = MARKER_DICT[abr_abbreviation]
        if ori.id == "BS_x7x_pKD46":
            assembly.abbreviation = f"{abr_abbreviation}7_pKD46"
            assembly.ori = "temperature sensitive pSC101"
        elif re.match("BS_x\dax", ori.id):
            assembly.abbreviation = abr_abbreviation + ori.id[-3:-1]
            assembly.ori = ORI_DICT[ori.id[-3]]
        else:
            assert re.match("BS_x\dx", ori.id) != None
            assembly.abbreviation = abr_abbreviation + ori.id[-2]
            assembly.ori = ORI_DICT[ori.id[-2]]
        assembly.id = f"BASIC_SEVA_{assembly.abbreviation}.10"
        assembly.description = f"BASIC SEVA vector containing {assembly.marker} resistance marker and {assembly.ori} origin of replication"
        assemblies.append(assembly)
build = bsb.BasicBuild(*assemblies)
PATH_TO_CSVS = Path.cwd().parents[0] / "csv_xlsx_files"
```

In [ ]:
```python
bsb.export_sequences_to_file(
    (assembly.return_part(
        description=assembly.description,
        name=assembly.id,
        id=seqrecord_hexdigest(assembly.return_part())
    ) for assembly in build.basic_assemblies),
    Path.cwd().parents[0] / "basicsynbio" / "parts_linkers" / "BASIC_SEVA_collection_v10.gb"
)
bsb.export_echo_assembly(
    build,
    PATH_TO_CSVS / "BASIC_SEVA_collection_v10_echo_scripts.zip",
    assemblies_per_clip=20,
    alternate_well=True,
)
bsb.pdf_instructions(
    build,
    "/home/hainesm6/github_repos/LondonBiofoundry/basicsynbio/pdfs/BASIC_SEVA_collection_v10_manual.pdf",
    assemblies_per_clip=20,
)
```

Out[ ]:
```
'/home/hainesm6/github_repos/LondonBiofoundry/basicsynbio/pdfs/BASIC_SEVA_collection_v10_manual.pdf'
```

In [ ]:
```python
# Calculate AbR marker mass to add to clip reaction
PJET_LEN = 2974
def approx_clip_mass(seq_length: int, ndigit: int =None) -> float:
    """Calculate the approximate mass of a BASIC part (ng) to add to a clip reaction."""
    return round(2.5*30e-6*(seq_length*607.4 + 157.9), ndigit)
abr_masses = {
    "Marker ID": [marker.id for marker in abr_markers],
    "mass (ng)": [
        approx_clip_mass(PJET_LEN + len(marker.seq)) for marker in abr_markers
    ],
}
abr_masses = pd.DataFrame(abr_masses)
abr_masses.to_csv(
    PATH_TO_CSVS / "AbR_marker_clip_masses.csv",
    index=False,
)
```

#### Results and discussion

- It has previously been reported that inhibition of protein synthesis dramatically increases the copy number of relaxed plasmids such as pBR322-ROP (Sambrook). This would lead to relatively higher expression of mScarlet compared to antibiotics that don't inhibit protein synthesis e.g. carbenicilin. This could explain why certain x9x constructs were burdensome.
- Lowering the concentration of tetracycline to 5 microgram/mL may improve the growth rate of BASIC_SEVA_5a5a.10, 5a7 and presumably 5a7_pKD46.10. All these plasmids replicate via a stringent mechanism and therefore cannot increase copy number if protein expression is blocked. This could explain why dropping the concentration improved the growth of these plasmids.

#### Aims and objectives for the cell/s below

- [x] Export csv file containing the following data for plasmids cultured at 37 degrees Celsius: name, marker, transformation well, overnight well.
- [x] Export a csv file containing data on BsaI digested constructs in the form: Assembly ID, Fragment 1 Length (bp), Fragment 2 Length (bp).
- [x] Generate a submission file for addgene for the collection.
- [X] Export `assembly.abbreviation` values to a spreadsheet.
- [x] What is the mean size of the plasmid in the collection.

In [ ]:
```python
def construct_picking_dict(
    overnight_wells: Tuple[str, ...],
    transformation_well: str,
    marker: str,
    name: str,
) -> Dict:
    """Return dictionary describing wells construct is picked into."""
    return [{
        "Overnight Well": overnight_well,
        "Transformation Well": transformation_well,
        "Marker": marker,
        "Name": name,
    } for overnight_well in overnight_wells]


PLATE_96 = Plate(size=96)


def plate_wells(
    number_of_wells: str,
    first_well_index: int,
    plate: Plate =PLATE_96,
):
    """slice wells of plate based on number of wells required and first well index."""
    return plate.wells[first_well_index:first_well_index + number_of_wells]


overnight_ind = 0
NUMBER_OF_WELLS_TO_PICK = 4
with open(PATH_TO_CSVS / "BASIC_SEVA_10_overnights.csv", "w", newline="") as csvf:
    fieldnames = ["Overnight Well", "Transformation Well", "Marker", "Name",]
    dictwriter = csv.DictWriter(csvf, fieldnames=fieldnames)
    dictwriter.writeheader()
    for ind, assembly in enumerate(build.basic_assemblies):
        if re.match("BASIC_SEVA_\d7_pKD46.10", assembly.id):
            continue
        elif re.match("BASIC_SEVA_\d[a-z]7_pKD46.10", assembly.id):
            continue
        dictwriter.writerows(construct_picking_dict(
            overnight_wells=plate_wells(
                NUMBER_OF_WELLS_TO_PICK,
                first_well_index=overnight_ind
            ),
            transformation_well=PLATE_96.wells[ind],
            marker=assembly.marker,
            name=assembly.id
        ))
        overnight_ind += NUMBER_OF_WELLS_TO_PICK
```

In [ ]:
```python
with open(PATH_TO_CSVS / "BASIC_SEVA_10_bsaI_digest.csv", "w", newline="") as csvf:
    fieldnames = ["Assembly ID", "Fragment 1 Length (bp)", "Fragment 2 Length (bp)"]
    dictwriter = csv.DictWriter(csvf, fieldnames=fieldnames)
    dictwriter.writeheader()
    for assembly_digest in bsb.build_digest(build):
        dictwriter.writerow({
            "Assembly ID": assembly_digest.assembly.id,
            "Fragment 1 Length (bp)": assembly_digest.product_lengths[0],
            "Fragment 2 Length (bp)": assembly_digest.product_lengths[1]
        })
```

In [ ]:
```python
addgene_template = pd.read_csv(PATH_TO_CSVS / "addgene_headers.csv")
ADDGENE_FIELDNAMES = tuple(name for name in addgene_template.columns)
rows = [
    {fieldname: "" for fieldname in ADDGENE_FIELDNAMES} for i in range(len(build.basic_assemblies))
]
for ind, assembly_digest in enumerate(bsb.build_digest(build)):
    rows[ind].update(
        {
            "Plasmid Name": assembly_digest.assembly.id,
            "Plasmid Type": "Encodes one insert",
            "Purpose": "BASIC DNA assembly vector",
            "Gene or insert Name": "mScarlet counter-selection cassette",
            "Insert Size": min(assembly_digest.product_lengths),
            "Species of gene or insert": "Synthetic",
            "Relevant Mutations": "Not applicable",
            "Backbone Name": assembly_digest.assembly.id,
            "Primary Vector Type": "Synthetic Biology",
            "Backbone size without insert": max(assembly_digest.product_lengths),
            "Cloning Method": "Unknown",
            "5-prime Sequencing Primer": "ggcggcggatttgtcctac",
            "3-prime Sequencing Primer": "ggtgagaatccaggggtcc",
            "Bacterial Resistance": "Spectinomycin" if assembly_digest.assembly.marker == "Streptomycin/Spectinomycin" else assembly_digest.assembly.marker,
            "high or low copy": "Low Copy",
            "Growth Temp": "30 C" if assembly_digest.assembly.ori == "temperature sensitive pSC101" else "37 C",
            "Growth Strain": "DH5alpha",
            "Hazardous": "No",
            "Patents or Licenses": "No",
            "Comments": "Cloning Method: BASIC DNA assembly",
            "Sequence:Full": str(assembly_digest.assembly.return_seqrec().seq)
        }
    )
with open(PATH_TO_CSVS / "BASIC_SEVA_10_addgene_batch_upload.csv", "w", newline="") as csvf:
    writer = csv.DictWriter(csvf, ADDGENE_FIELDNAMES)
    writer.writeheader()
    writer.writerows(rows)
```

In [ ]:
```python
abbreviation_df = pd.DataFrame(
    {
        "BASIC SEVA ID": [seqrecord_hexdigest(assembly.return_part()) for assembly in build.basic_assemblies],
        "Abbreviations": [assembly.abbreviation for assembly in build.basic_assemblies],
    }
)
abbreviation_df.to_csv(
    PATH_TO_CSVS / "BASIC_SEVA_10_abbreviations.csv",
    index=False
)
```

In [ ]:
```python
print(f"Mean plasmid size: {np.mean(np.array([len(assembly.return_part()) for assembly in build.basic_assemblies]))}")
```

```
Mean plasmid size: 4025.4
```

#### Aims/objectives for cell/s below

- [x] Export sequence flanked by *i*P and *i*S regions in oris for use by cmatch to validate assemblies.

In [ ]:
```python
core_ori_seqs = [ori.basic_slice() for ori in oris]
for core_ori_seq in core_ori_seqs:
    SeqIO.write(
        core_ori_seq,
        PATH_TO_SEQS / "genbank_files" / "BASIC_SEVA_collection" / "misc_seqs" / f"{core_ori_seq.id}_core_seq.gb",
        "genbank"
    )
```
