## Supplementary_Data for "basicsynbio and the BASIC SEVA collection: Software and vectors for an established DNA assembly method": BASIC_SEVA_collection_v10_manual.pdf

### Protocol for BasicBuild 22-12-2021

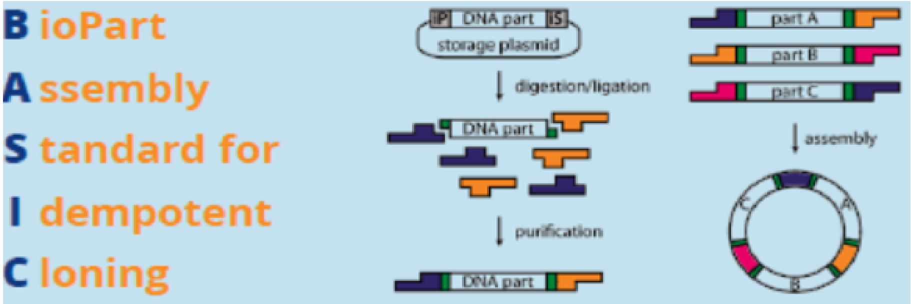


### Materials

| Item | Order number (if applicable) |
|---|---|
| PCR machine | |
| Water bath (42°C) for transformation | |
| Magnetic plate | Ambion AM10050 (Thermo) |
| Benchtop centrifuge | |
| Microplate centrifuge | |
| Vortex | |
| 96 well U-Bottom Falcon plate | Falcon 351177 (Thermo) |
| Microcentrifuge tubes | |
| PCR tubes | |
| Brooks Life Sciences PCR Foil Seals | 4ti-0550 |
| Pipettes/tips 10 and 200 µl | |
| Agencourt AMPure XP SPRI paramagnetic beads | A6388 (Beckman Coulter) |
| ddH20 | |
| 70% EtOH | |
| Biolegio BASIC linkers | |
| NEB BsaI-HF v2 enzyme (R3733) 20 U/µl | R3733 (NEB) |
| Promega T4 ligase (M1801) 1-3U/µl | M1801 (Promega) |
| [optional] 10x Assembly Buffer: 0.2 M Tris:HCl (pH 8.0), 0.1 M MgCl2, 0.5 M KCl | |
| Chemically competent cells (DH5alpha, 1x109 CFU/µg pUC19 | C2987I (NEB) or equivalent |
| SOC media | |
| Petri dishes | |
| LB-Agar + antibiotic/s | |

### Protocol for BasicBuild 22-12-2021

### Method

#### Preperation of BASIC linkers and parts

BASIC linkers can be ordered from Biolegio and will be delivered in a lyophilized format along with linker annealing buffer.

1. Spin down plates containing lyophilized linkers.
2. Add 150 µl of linker annealing buffer to each well, seal the plate with a PCR foil and incubate for 1 hour at room temperate.
3. Vortex the plate and collect liquid via centrifugation.
4. Conduct the following incubation in a thermocycler:

| Temperature (°C) | Time | |
|---|---|---|
| 95 | 2 min | |
| 94 (-1 °C/cycle) | 40 seconds | x70 cycles |
| 4 | Hold | |

5. Collect the liquid in wells via centrifugation.
6. Linkers are ready to use or can be stored at -20°C until required.

#### Clips Reaction

Below contains a table with each clip needed for the assemblies of the build.

| Clip Index | Prefix ID | Part Name | Part mass for 30 µL clip reaction (ng) | Suffix ID | Total ass emblies | Assembly indexes | Clip plate mapping |
|---|---|---|---|---|---|---|---|
| 1 | LMP-P | BS_1xx | 57 | BSEVA_L1-S | 5 | [1, 2, 3, 4, 5] | nan |
| 2 | BSEVA_L1-P | BS_x5ax | 237 | LMS-S | 6 | [1, 6, 11, 16, 21, 26] | nan |
| 3 | LMS-P | B407_J23119-RBS34-mScarl | 139 | LMP-S | 12 | [1, 2, 6, 7, 11, 12, 16, 17, 21, 22, 26, 27] | nan |
| 4 | BSEVA_L1-P | BS_x6x | 100 | LMS-S | 6 | [2, 7, 12, 17, 22, 27] | nan |
| 5 | BSEVA_L1-P | BS_x7x | 134 | LMS-S | 6 | [3, 8, 13, 18, 23, 28] | nan |
| 6 | LMS-P | B408_J23119-RBS-A12-mSc | 181 | LMP-S | 12 | [3, 5, 8, 10, 13, 15, 18, 20, 23, 25, 28, 30] | nan |
| 7 | BSEVA_L1-P | BS_x9x | 131 | LMS-S | 6 | [4, 9, 14, 19, 24, 29] | nan |

| | | | | | | | |
|---|---|---|---|---|---|---|---|
| 8 | LMS-P | B405_J23106-RBS34-mScarl | 169 | LMP-S | 6 | [4, 9, 14, 19, 24, 29] | nan |
| 9 | BSEVA_L1-P | BASIC_construct_BS_x7x_ | 137 | LMS-S | 6 | [5, 10, 15, 20, 25, 30] | nan |
| 10 | LMP-P | BS_2xx | 51 | BSEVA_L1-S | 5 | [6, 7, 8, 9, 10] | nan |
| 11 | LMP-P | BS_3xx | 45 | BSEVA_L1-S | 5 | [11, 12, 13, 14, 15] | nan |
| 12 | LMP-P | BS_4xx | 55 | BSEVA_L1-S | 5 | [16, 17, 18, 19, 20] | nan |
| 13 | LMP-P | BS_5axx | 67 | BSEVA_L1-S | 5 | [21, 22, 23, 24, 25] | nan |
| 14 | LMP-P | BS_6xx | 46 | BSEVA_L1-S | 5 | [26, 27, 28, 29, 30] | nan |

Prepare a Master mix for clip reactions, the below table provides the required components for the master mix with sufficient quantities for all clip reactions.

| Component | Volume per clip (µL) |
|---|---|
| Promega T4 DNA Ligase 10x Buffer | 45.0 |
| Water | 232.5 |
| NEB BsaI-HFv2 | 15.0 |
| Promega T4 DNA Ligase | 7.5 |

For each Clip reaction, setup 1 PCR tube with 30 µl total volume:

Dispense 20 uL master mix, 1 µl of each prefix and suffix Linker, 1 µl of part (or more depending on concentration) into a PCR tube and make up to 30 µl with water.

After mixing, tubes are placed in a PCR machine running the following programme:

| Temperature (°C) | Time | |
|---|---|---|
| 37 | 2 min | x20 cycles |
| 20 | 1 min | |
| 60 | 5 min | |

#### Magbead purification

Prepare fresh 70% EtOH (0.5 ml per BASIC reaction) and bring magnetic beads (AmpureXP or Ampliclean) stored at 4°C back into homogeneous mix by shaking thoroughly.

1. Add 54 µl of magnetic beads into 96 well Falcon plate (one well per BASIC reaction) and add the 30 µl BASIC linker ligation from the PCR machine step, mix by pipetting 10 times.
2. Wait 5 min to allow DNA binding to magbeads.
3. Place Falcon plate on magnetic stand and wait for rings to form and solution to clear.
4. Aspirate most of the solution with a 200 µL pipette set to 80 µL.
5. Add 190 µl 70% EtOH to each well and wait 30 s.

### Protocol for BasicBuild 22-12-2021

6 Remove solution from each well (pipette set to 200 µl volume)

7 Add 190 µl 70% EtOH to each well and wait 30 s.

8 Remove solution from each well (pipette set to 200 µl volume)

9 Leave the plate to dry for 1-2 min.

10 Remove Falcon plate from magnet and resuspend magbeads in 32 µl dH20.

11 Wait 1 min for DNA to elute.

12 Place Falcon plate back on magnetic stand and allow ring to form and solution to clear.

13 Transfer 30 µl of purified clip reaction into a clean microcentrifuge tube or well and store at at -20°C for up to 1 month.

#### Assembly reaction

Below contains a table with each BASIC assembly within your build

| Assembly Index | Assembly ID | Clip indexes | Assembly plate mapping |
|---|---|---|---|
| 1 | BASIC_SEVA_15a.10 | [1, 2, 3] | nan |
| 2 | BASIC_SEVA_16.10 | [1, 4, 3] | nan |
| 3 | BASIC_SEVA_17.10 | [1, 5, 6] | nan |
| 4 | BASIC_SEVA_19.10 | [1, 7, 8] | nan |
| 5 | BASIC_SEVA_17_pKD46.10 | [1, 9, 6] | nan |
| 6 | BASIC_SEVA_25a.10 | [10, 2, 3] | nan |
| 7 | BASIC_SEVA_26.10 | [10, 4, 3] | nan |
| 8 | BASIC_SEVA_27.10 | [10, 5, 6] | nan |
| 9 | BASIC_SEVA_29.10 | [10, 7, 8] | nan |
| 10 | BASIC_SEVA_27_pKD46.10 | [10, 9, 6] | nan |
| 11 | BASIC_SEVA_35a.10 | [11, 2, 3] | nan |
| 12 | BASIC_SEVA_36.10 | [11, 4, 3] | nan |
| 13 | BASIC_SEVA_37.10 | [11, 5, 6] | nan |
| 14 | BASIC_SEVA_39.10 | [11, 7, 8] | nan |
| 15 | BASIC_SEVA_37_pKD46.10 | [11, 9, 6] | nan |
| 16 | BASIC_SEVA_45a.10 | [12, 2, 3] | nan |
| 17 | BASIC_SEVA_46.10 | [12, 4, 3] | nan |
| 18 | BASIC_SEVA_47.10 | [12, 5, 6] | nan |
| 19 | BASIC_SEVA_49.10 | [12, 7, 8] | nan |
| 20 | BASIC_SEVA_47_pKD46.10 | [12, 9, 6] | nan |
| 21 | BASIC_SEVA_5a5a.10 | [13, 2, 3] | nan |
| 22 | BASIC_SEVA_5a6.10 | [13, 4, 3] | nan |
| 23 | BASIC_SEVA_5a7.10 | [13, 5, 6] | nan |
| 24 | BASIC_SEVA_5a9.10 | [13, 7, 8] | nan |
| 25 | BASIC_SEVA_5a7_pKD46.10 | [13, 9, 6] | nan |

| | | | |
|---|---|---|---|
| 26 | BASIC_SEVA_65a.10 | [14, 2, 3] | nan |
| 27 | BASIC_SEVA_66.10 | [14, 4, 3] | nan |
| 28 | BASIC_SEVA_67.10 | [14, 5, 6] | nan |
| 29 | BASIC_SEVA_69.10 | [14, 7, 8] | nan |
| 30 | BASIC_SEVA_67_pKD46.10 | [14, 9, 6] | nan |

For each BASIC assembly, combine the required purified clip reactions in a final volume of 10 µl in 1x Assembly or NEB CutSmart buffer as follows:

| Reagent | Volume |
|---|---|
| 10x Assembly Buffer (or 10x NEB CutSmart) | 1 µl |
| Each purified clip reaction required for the assembly | 1 µl for each |
| dH20 | Top up to 10 µl total volume |

Run assembly reaction in PCR machine with following programme

| Temperature (°C) | Time |
|---|---|
| 50 | 45 min |
| 4 | Hold |

#### Transformation

Use 50 µl of chemically competent cells DH5alpha with high transformation efficiency (109 CFU/µg pUC19, for instance NEB C2987I) to transform 5 µl of each BASIC assembly:

1. Chemically competent cells are stored at -80°C.
2. Thaw competent cells on ice (takes 5-10 min); 50 µl per BASIC assembly to be transformed.
3. Cool 5 µl of BASIC DNA assembly in 1.5 ml microcentrifuge tube on ice.
4. Add 50 µl of competent cells to each precooled 5 µl BASIC reaction.
5. Incubate on ice for 20 min.
6. Apply heat shock in 42°C water bath for 45s and place back on ice for 2 min.
7. Add 200 µl SOC medium to each tube and incubate shaking at 37°C for 1h recovery.
8. Spot or plate cells on agar plates with appropriate antibiotics. Depending on number of parts assembled and transformation efficiency 2-250 µl might be spotted or plated.
9. Incubate agar plates at 37°C overnight, next day pick colony for assay or miniprep.
